# Automatic diagnostics of electroencephalography pathology based on multi-domain feature fusion

**DOI:** 10.1101/2024.09.01.610707

**Authors:** Shimiao Chen, Dong Huang, Xinyue Liu, Jianjun Chen, Xiangzeng Kong, Tingting Zhang

**Author notes:** These authors contributed equally to this work (Shimiao Chen, Dong Huang).

## Abstract

Electroencephalography (EEG) serves as a practical auxiliary tool deployed to diagnose diverse brain-related disorders owing to its exceptional temporal resolution, non-invasive characteristics, and cost-effectiveness. In recent years, with the advancement of machine learning, automated EEG pathology diagnostics methods have flourished. However, most existing methods usually neglect the crucial spatial correlations in multi-channel EEG signals and the potential complementary information among different domain features, both of which are keys to improving discrimination performance. In addition, latent redundant and irrelevant features may cause overfitting, increased model complexity, and other issues. In response, we propose a novel feature-based framework designed to improve the diagnostic accuracy of multi-channel EEG pathology. This framework first applies a multi-resolution decomposition technique and a statistical feature extractor to construct a salient time-frequency feature space. Then, spatial distribution information is channel-wise extracted from this space to fuse with time-frequency features, thereby leveraging their complementarity to the fullest extent. Furthermore, to eliminate the redundancy and irrelevancy, a two-step dimension reduction strategy, including a lightweight multi-view time-frequency feature aggregation and a non-parametric statistical significance analysis, is devised to pick out the features with stronger discriminative ability. Comprehensive examinations of the Temple University Hospital Abnormal EEG Corpus V. 2.0.0 demonstrate that our proposal outperforms state-of-the-art methods, highlighting its significant potential in clinically automated EEG abnormality detection.

## Introduction

Electroencephalography (EEG) is a non-invasive neuroimaging technology that monitors and records the bioelectric signals arising spontaneously from the electrical activity of brain neurons [1–3]. Due to its portability, cost-effectiveness, high time resolution, and ease of implementation, EEG is widely used in clinical diagnosis of various neurological disorders, e.g., seizures [4–6], depressive disorder [7, 8], and Parkinson’s disease [9, 10], where the task of distinguishing between non-pathological and pathological EEG patterns at outset is forefront [11]. Based on this classification, further investigations or the prescription of medication can be made. Traditionally, proficient clinicians or neurologists meticulously scrutinize 20-30 minute brainwave recordings to identify subtle changes in frequency or amplitude that may convey crucial physiological and pathological information for EEG detection [12–14]. Not only is this process time-consuming and labor-intensive, but also requires years of training for physicians to obtain board certification, resulting in a shortage of qualified neurologists and professional EEG analyzers [15–17]. In addition, experts usually adopt a complex decision tree to analyze and categorize these signals [18], which would give rise to inter-rater disagreements. Therefore, the development of an automated EEG classification methodology without human intervention is essential to deliver accessible and reliable clinical EEG diagnosis services for hospitals and medical centers.

In recent years, machine learning has attracted widespread attention in the research community of pathological EEG diagnosis, which can be broadly categorized into deep learning and feature-based approaches. Deep learning approaches, including convolutional neural networks (CNNs) [14], long short-term memories (LSTMs) [3], and temporal convolutional networks [11], can automatically extract and classify features from raw data. Adopting increasingly complex frameworks can yield performance improvements, since neural network performance exhibits a power-law correlation with model complexity and training data [13, 19]. Unfortunately, not only is this improvement marginal, but the excessively large architecture also causes various other problems, such as gradient vanishing or gradient explosion [14, 16].

As promising alternatives, feature-based approaches have several advantages, such as relatively low computational burden and notable performance enhancements within a simple structure. Adhere to a ”feature engineering→classification” structure, it initially learns various informative EEG features, selects the optimal feature subsets from the constructed feature space, and then feeds these features into the classifier for discerning pathology EEG. However, the intricate nonlinear dynamics, non-stationarity, weak signal strength, and susceptibility to noise and artifacts inherent in EEG make effective feature engineering difficult and potentially damage the EEG classification performance [20, 21]. Therefore, adopting a suitable and effective feature analysis technique is the key to the success of feature-based approaches.

To date, a large number of studies have focused on temporal and frequency domain feature analysis [15, 22]. These methods can easily and rapidly learn linear features and interpretable representations from various single domains. Nevertheless, they usually ignore other domain-specific features, leading to insufficient characterization of the low signal-to-noise ratio (SNR) and nonlinear EEG signals. In particular, time-domain feature extraction fails to consider energy distribution and spatial relationships among various brain regions, while frequency-domain analysis lacks time-varying statistical properties and spatial features [23]. Diverging from these two domain techniques, joint time-frequency feature extraction has been propelled into the spotlight by its greater adaptability, sensitivity to transient changes, and ability to capture frequency components changing over time. Representative methodologies in this domain include Wavelet Packet Decomposition (WPD) [20, 24], Short-time Fourier transform (STFT) [16, 25], and Discrete Wavelet Transform (DWT) [2, 11].

After feature representation, an efficient and accurate classifier is imperative for assigning the appropriate label to each test EEG sample. Commonly utilized classifiers encompass Support Vector Machine (SVM) [26, 27], *K*-Nearest Neighbors (KNN) [2, 28], and Gradient Boosting Decision Trees (GBDTs). Notably, GBDTs including but not limited to Categorical Boosting (CatBoost) [29], Extreme Gradient Boosting (XGBoost) [20], and Light Gradient Boosting Machine (LightGBM) [16] have been extensively utilized in EEG classification, as they are more excellent and more robust compared to conventional single classifiers.

Although significant progress in the EEG detection domain made by machine learning, there are several limitations that should be noticed and addressed. Firstly, the majority of existing methods primarily focus on feature extraction but neglect the important feature selection, resulting in a substantial number of retained redundant and irrelevant information, detrimentally affecting the EEG classification performance. Secondly, few works have explored spatial domain information in multi-channel EEG signals, limiting the ceiling of diagnostics precision. Thirdly, the complementarity among hierarchical features from temporal, spectral, and spatial domains has been neglected in EEG pathology detection, even though it has proven to be effective in many other EEG analyses [1, 30]. Thus, how to learn high-quality information representation to improve automated EEG diagnosis remains a persistent concern.

To overcome the aforementioned limitations, this work proposes a novel feature-based framework for multi-channel EEG detection. Specifically, this framework comprises three main components: (i) To take full advantage of the complementarity among different domain features, we introduced a multi-feature learning mechanism, consisting of a time-frequency feature extractor and a spatial feature extractor. The former is devised to learn salient time-frequency features with the help of a multi-resolution DWT decomposition mechanism and a statistical feature extractor. In addition, it is important to note that, unlike traditional techniques, the latter mines subtle spatial features from time-frequency information, thereby enhancing the performance of subsequent tasks. (ii) Considering the potential risks posed by high-dimensional features, a two-step dimension reduction is used to wipe out unnecessary features. The first step is a multi-view aggregation applied to the extracted time-frequency features which are subsequently combined with spatial features, while the second step is a statistical significance analysis to validate the fused results. (iii) Lastly, the optimal feature set is input into several different ensemble learning classifiers to categorize EEGs as either normal or abnormal. Extensive experiments on the publicly accessible EEG database demonstrate that the proposed methodology surpasses competitive methodologies. Additionally, ablation experiments further confirm the exceptional efficacy of the devised feature analysis technique in enhancing information comprehensiveness and refinement. In a nutshell, the major contributions of this study are listed as follows:

1. We introduce a novel multi-domain feature fusion strategy that densely integrates EEG features across temporal, spectral, and spatial domains to provide comprehensive information representation. Additionally, the spatial information is derived from the denoised time-frequency information instead of raw EEG signals, improving the performance of the EEG detection system.
2. We propose an innovative two-step strategy to reduce the dimension of the constructed feature space and pick out the most discriminative information, thereby enhancing the capability of feature expression.
3. Comprehensive experiments on a real-world EEG dataset showcase our method’s superiority over state-of-the-art baselines. Moreover, ablation studies substantiate the efficacy of the proposed multi-view feature aggregation and spatial information extraction.

The subsequent sections of this paper are organized in the following manner. In Section, we provide a comprehensive overview of deep learning methodologies and feature-based methodologies proposed in recent years. Section outlines the EEG benchmark dataset used for experiments and elaborates on the designed automatic diagnostics methodology of EEG pathology. Section discusses the comparative experiments, and Section presents a conclusive summary and sets the course for future research.

## Related Work

Over the past few years, numerous approaches have been proposed in the literature, aimed at effectively and automatically discerning pathological EEG signals from their normal counterparts. They can be roughly divided into two categories: deep learning approaches and feature-based approaches. This a concise yet comprehensive overview of recent works associated with these two categories.

### Deep Learning Methods

In recent years, substantial efforts have been made to tackle the challenge in general EEG pathology classification through the assistance of deep learning approaches. For instance, Schirrmeister et al. [31] designed a 4-layer deep ConvNet architecture namely BD-Deep4 to identify anomalous events in EEGs, achieving an accuracy of 85.42%. Roy et al. [19] put forward a deep 1D convolutional gated recurrent neural network, i.e., ChronoNet, resulting in 86.57 % classification accuracy. In [14], the authors introduced a novel deep one-dimensional CNN model for the discrimination of two EEG patterns. Moreover, there has been an increasing interest in employing transfer learning and hybrid deep learning techniques for the automatic categorization of EEGs into normal or abnormal. Amin et al. [32] deployed a pre-trained AlexNet model with the last layer replaced by an SVM classifier to conduct EEG pattern recognition, reporting an accuracy of 87.32%. Also, in [22], an AlexNet pre-trained in a non-disclosed database was used to extract features from cropped data, accompanied by a Multilayer Perceptron (MLP) for classification. Beyond that, Khan et al. [3] developed a novel hybrid model that integrates CNN-based feature extraction and LSTM-based classification, resulting in 86.23% accuracy. Likewise, Albaqami et al. [33] combined customized WaveNet and LSTM sub-models to differentiate EEG signals and obtain an accuracy rate of 88.76%.

To comprehensively explore the nature of deep learning approaches, recent studies have undertaken a substantial amount of comparative analysis. In [11], various CNN-based models were systematically compared and analyzed on Temple University Hospital (TUH) dataset. The optimized temporal convolutional network with 456,502 trainable parameters demonstrated superior performance in classifying pathological and non-pathological signals. More recently, Kiessner et al. [17] conducted another holistic examination to evaluate the EEG decoding performance of various deep neural networks on an extended EEG dataset, which is five times larger than the TUH dataset. The outcomes showcased that the most complex model with over four hundred thousand parameters achieved the best performance accuracy of 86.59%.

In summary, even though deep learning approaches can yield marginal improvements in the automatic pathological EEG identification, this is at the cost of a more complex model architecture, a greater need for labeled training samples, and increased computational time and storage requirements [16, 17, 31]. Moreover, collecting well-labeled datasets is arduous and prone to a relatively low inter-rater agreement, while designing a complex yet high-performing end-to-end structure is difficult [11, 34]. Furthermore, the spatial correlation embedded in multi-channel EEG data is another essential yet under-explored factor for accurate EEG detection.

### Feature-based Methods

In recent years, feature-based approaches have gained prominence in EEG pathology diagnosis, due to lower hardware requirements, simpler model architecture, adaptability in learning meaningful features, and other advantages [2, 25]. These approaches primarily follow two stages: feature engineering and classification. Notably, the former including feature extraction and feature selection is critical for boosting EEG decoding performance due to the fact that the classification accuracy highly relies on the learned feature.

Up to now, numerous feature extraction methodologies have been proposed to cope with EEG binary classification, which can be broadly grouped into single-domain, dual-domain, and multi-domain techniques. Representative single-domain analysis techniques are temporal domain analysis methods and frequency domain analysis methods. For instance, various spectral features are captured from channel-, segment-, and EEG-level to detect pathological slowing in EEG signals [15]. The dual-resolution analysis techniques, especially the time-frequency analysis, have garnered increasing attention in discerning EEG patterns owing to their ability to mine complementary information among two distinct domains. For example, Cisotto et al. [13] computed eleven well-established time-frequency domain features in each frame of each EEG channel to distinguish normal and abnormal EEGs. Sharma et al. [12] extracted fuzzy entropy, logarithmic of the squared norm, and fractal dimension feature from wavelet sub-bands, obtaining an accuracy rate of 79.34% with the assistance of the SVM classifier. Similarly, Singh et al. [25] converted brain signals into images via STFT and achieved a classification accuracy of 88.04%. In [2], a hypercube-based feature extractor coupled with DWT was used to decompose signals into a series of physically meaningful narrow-band signals, in conjunction with Neighborhood Component Analysis-based feature selection and the KNN classifier, achieving the accuracy of 87.68%. Along a similar line, Gemein et al. [11] devised EEG pathology diagnosis methods based on DWT. More recently, Zhong et al. [24] implemented WPD and various ensemble learning classifiers to classify EEG data, which achieved a state-of-the-art accuracy of 89.13%.

It is evident from the above research that feature-based approaches possess tremendous potential in boosting the binary classification performance of multi-channel EEG records. However, few works took the subtle spatial information into account and made use of the mutual complementarity between time, frequency, and spatial domains, causing a decline in classification performance. Moreover, the redundant and irrelevant features embedded in the extracted features, which may cause overfitting and increased computational costs, pose another challenge for EEG pathology detection. Therefore, we put forward a novel feature-based traditional machine learning framework to distinguish normal and abnormal EEG, which integrates a multi-domain feature fusion and a two-step dimension reduction to provide refined and comprehensive information representation.

## Materials and Methods

In this section, we develop an innovative framework for automatically detecting abnormal EEG patterns from normal ones. As shown in Fig 1, its pipeline includes the following primary phases: Firstly, a preprocessing phase is performed on raw EEG to guarantee data uniformity. The second phase is capturing and fusing features from multiple domains, aimed at enhancing the representation capabilities of features. Subsequently, a two-step dimension reduction strategy is implemented to remove redundant and non-informative information. The resulting features are fed into three distinct traditional machine-learning algorithms for EEG classification. Detailed descriptions of each phase are provided in the subsequent subsections.

**Figure 1.**
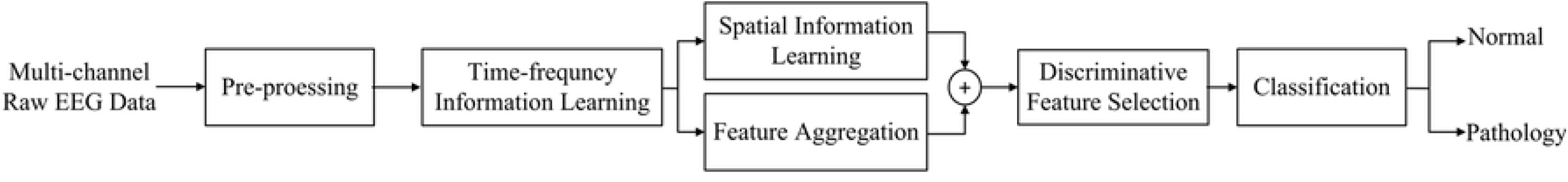
Block diagram of the proposed abnormal EEG signal de

### Data Description and Preprocessing

To verify the practicality of the proposed method, a large-scale EEG dataset, known as the TUH Abnormal EEG Corpus [35], was employed. This dataset, continuously updated and currently at version 2.0.0, is the most comprehensive open-source EEG benchmark for evaluating abnormal EEG detection systems. The scalp EEG recordings were gathered from 2,329 distinct patients, spanning ages from 7 days to 96 years, covering diverse diagnoses including but not limited to epilepsy, sleep disorders, and brain injuries. All EEG recordings were collected using the international 10-20 sensor placement system at a sampling frequency of 250 Hz or higher, lasting for at least 15 minutes. 1,521 instances were manually marked as normal, and 1,472 were labeled as pathological. The corpus was split into two exclusive subsets, with 70% allocated for training and the remaining 30% for testing. The specific details, including data division, gender distribution, and so on, are provided in Table 1.

**Table 1.** Detailed description of the TUH Abnormal EEG dataset utilized in this study.

| Description |  | Training |  |  | Evaluation |  |  |
| --- | --- | --- | --- | --- | --- | --- | --- |
|  |  | Female | Male | Total | Female | Male | Total |
| Normal | Recordings | 768 | 603 | 1371 | 85 | 65 | 150 |
|  | Patients | 691 | 546 | 1237 | 84 | 64 | 148 |
| Pathology | Recordings | 679 | 667 | 1346 | 63 | 63 | 126 |
|  | Patients | 454 | 439 | 893 | 51 | 54 | 105 |

In the EEG pathology detection system, the preprocessing step is usually taken to ensure consistency and reduce useless information, facilitating more accurate and reliable results [13, 33]. Therefore, we apply a three-stage preprocessing process: channel selection, downsampling, and signal segmentation, as shown in Fig 2. Firstly, to reduce unnecessary information and maintain data consistency, we selected the same 21 channels conforming to the 10-20 International montage (refer to Fig 3) in all recordings. Secondly, the EEG samples were downsampled to a frequency rate of 250 Hz to mitigate large transients’ impact and accelerate the computation, in accordance with the work [19]. Lastly, EEG recordings were segmented channel-wise into 100 non-overlapping partitions using a 5 s sliding window. The extra recordings were abandoned.

**Figure 2.**
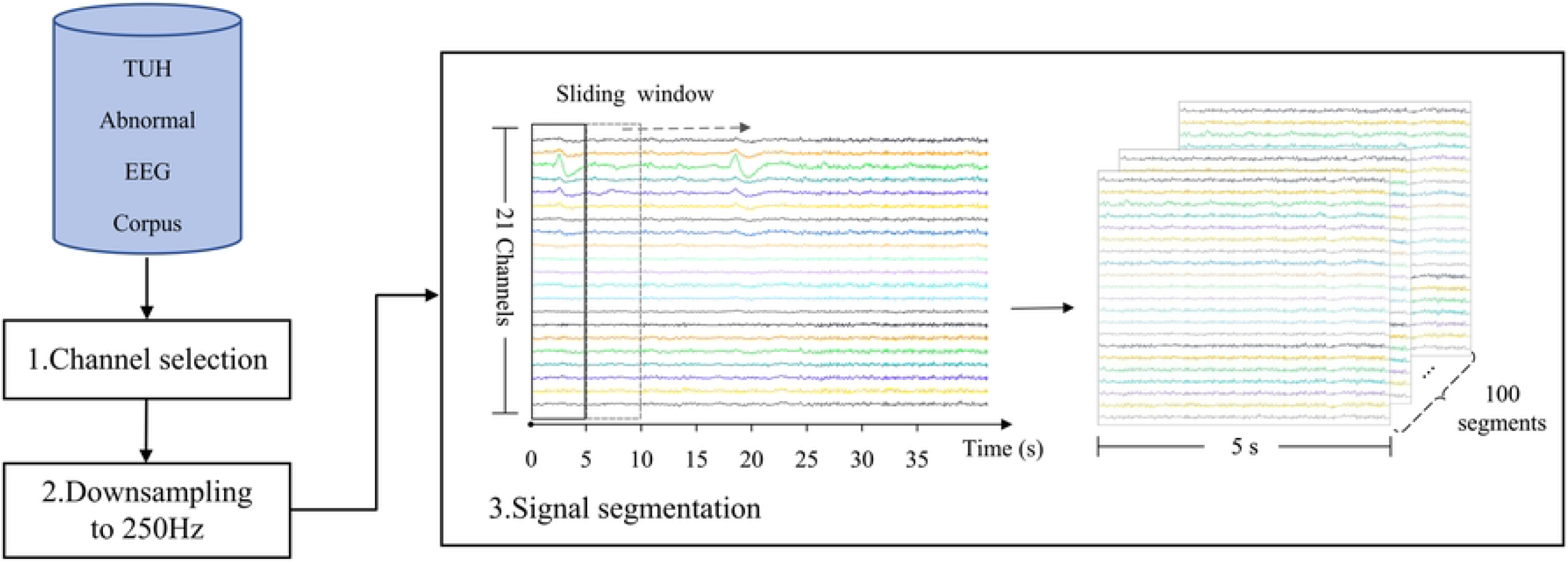
Flowchart of EEG preprocessing.

**Figure 3.**
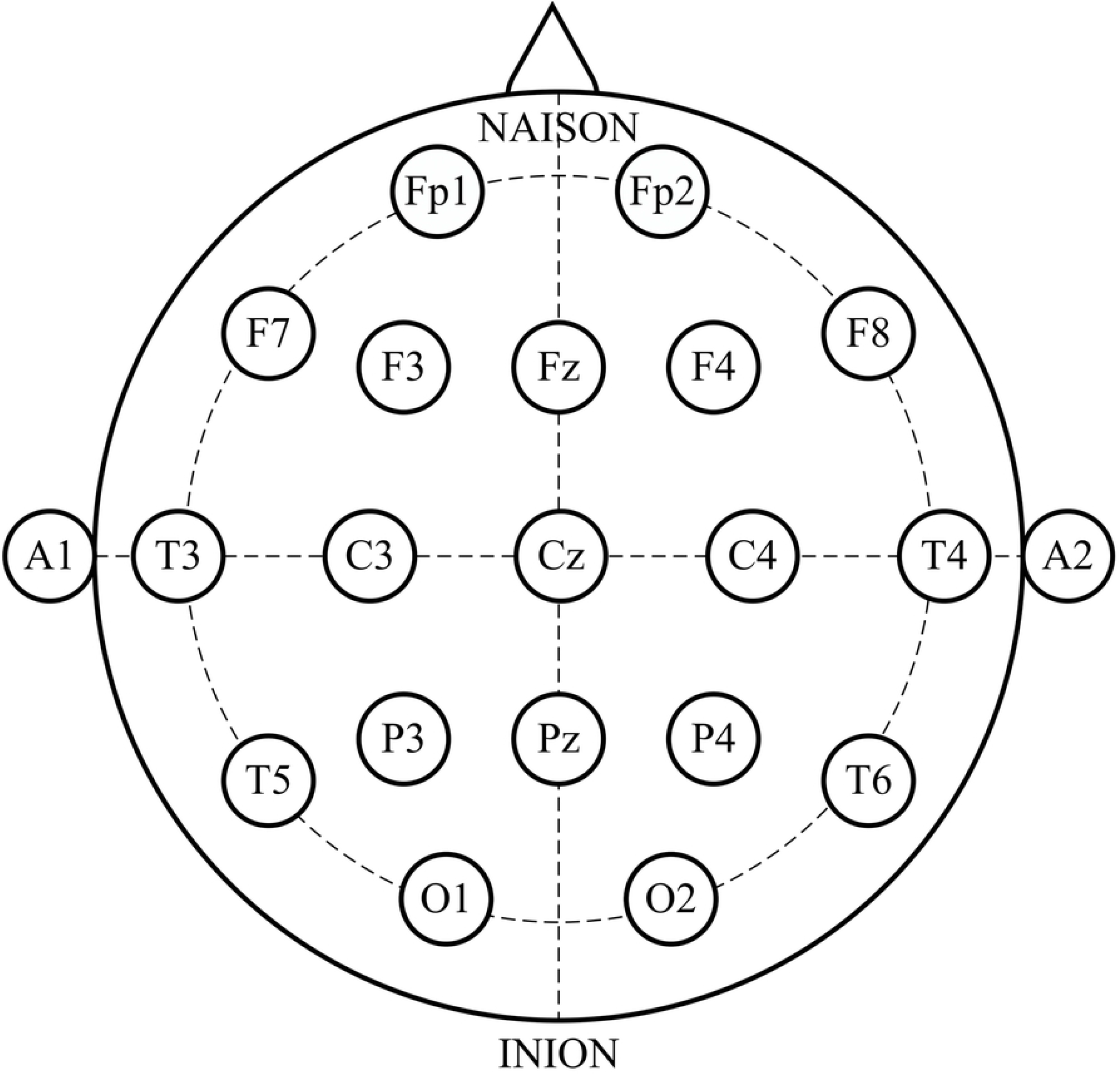
Distribution of the 21 EEG electrodes according to 10-2

### Multi-domain Feature Extraction

EEG signals encompass intricate and abundant information in time, frequency, and spatial domains [30]. Relying solely on extracting features from a single domain may fall short of capturing the complex nature of this signal. Hence, to overcome this limitation, we attempt to analyze EEG from multiple perspectives simultaneously in this work.

### Multi-resolution Dual-domain Analysis

Recently, plenty of temporal-frequency domain analysis techniques have been introduced to enhance the accuracy in discerning abnormal brain signals. These dual-domain techniques excel in revealing intricate details concerning the amplitude and phase variations of different frequency components over time. A common technique is STFT, which can capture the signal’s frequency content and time localization information over a defined sliding window [16]. However, its pre-fixed analysis window renders it non-adaptive in multi-resolution dual-domain analysis. Conversely, the wavelet transform techniques, including DWT and WPD, can realize time subdivisions at high frequencies while frequency subdivisions at low frequencies by stretching and translating wavelet functions. However, DWT typically splits high-frequency bands into more subtle subbands, while WPD processes both low- and high-frequency components and generates more subsequences, exacerbating the problem of redundant and irrelevant information as well as higher time complexity [26, 28]. This is the rationale behind the utilization of DWT.

DWT can recursively decompose EEG signals into multi-resolution frequency sub-bands at finite layers, as illustrated in Fig 4. At *j*-layer (*j* = 1, 2, …, *J*), the input is broken into high- and low-frequency sub-bands of the same scale respectively through concurrent convolution with a high-pass filter *h*_*j*_ and a low-pass filter *g*_*j*_. Following this, the intermediate sub-bands undergo 1/2 downsampling to generate an approximate component *A*_*j*_ (Eq (1)) and a detail component *D*_*j*_ (Eq (2)). The former describes the signal’s long-term trend and reflects overall identity, while the latter captures short-term trends and subtle nuances in sub-bands. Notably, the approximate component *A*_*j*_ typically serves as the input for the (*j* + 1)-layer. This process is iterated until reaching the final decomposition layer *J*.

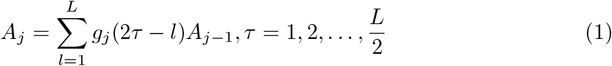

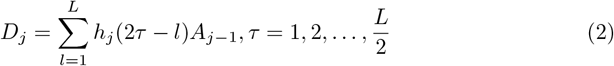

where *L* stands for the length of *A*_*j*−1_ after *j*− 1 decompositions and *τ* stands for the scale.

**Figure 4.**
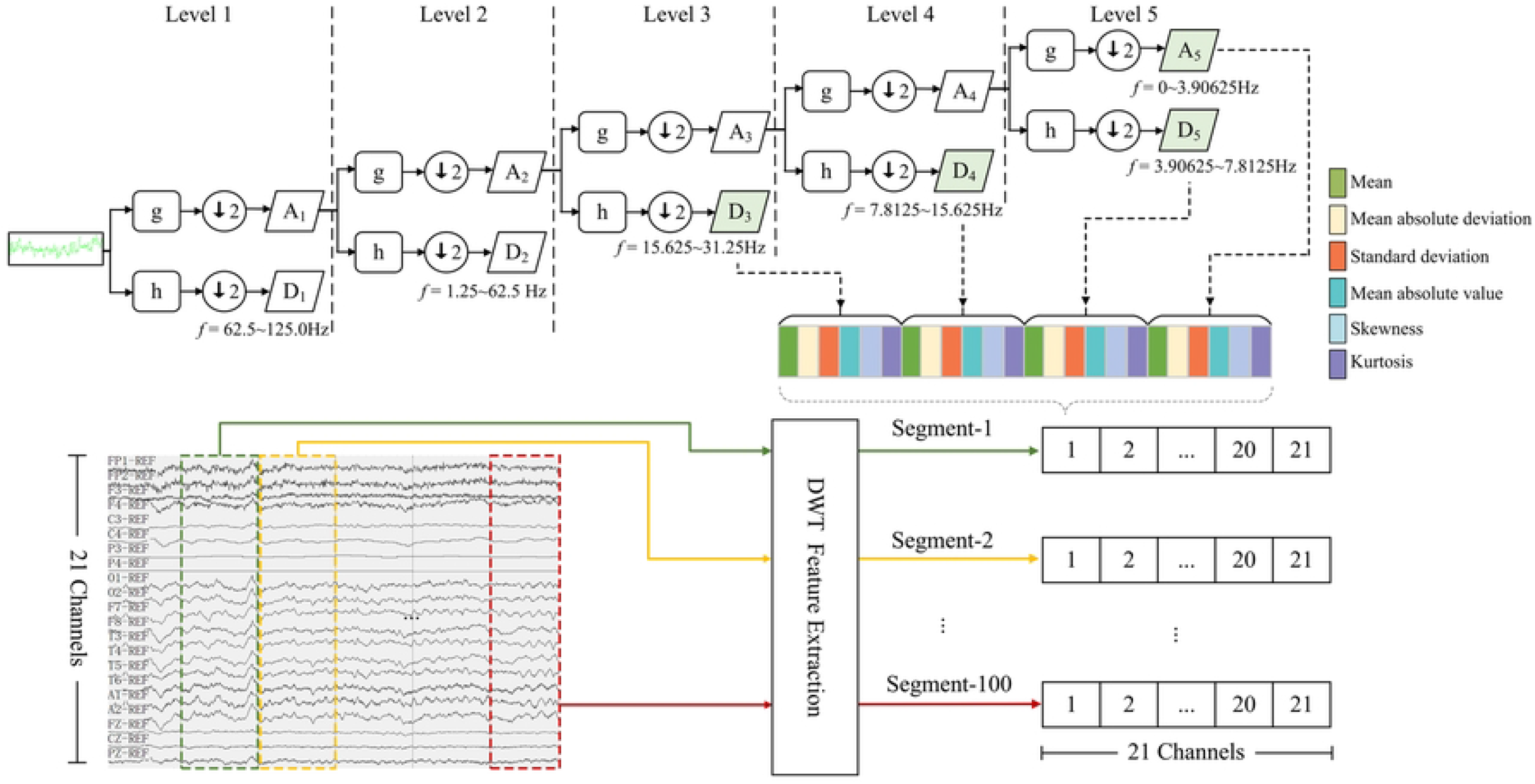
Time-frequency feature extraction on raw EEG signals

The decomposition level and the applied mother wavelet significantly impact the quality of EEG decomposition, since they determine the structure of the filter bank and the resulting frequency components. According to [29, 36], brainwave rhythms are categorized basically into five distinct frequency bands: Delta, Theta, Alpha, Beta, and Gamma (above 30 Hz); of these, the first four bands have been widely used to study brain function. As an illustration, the features associated with epileptic seizure predominantly appear in the frequency spectrum below 30 Hz [28], while the study [8] reveals that distinct hemisphere asymmetry differences between affected and unaffected subjects can be investigated by assessing the relative signal power in four bands.

Therefore, this work decomposed 250 Hz EEG signals using a 5-level DWT with Symlets wavelet of order 6 (sym6), whose orthogonality is particularly suitable for distinguishing abnormal EEG patterns [37]. After wiping out irrelevant components, *D*_3_ (15.625-31.25 Hz), *D*_4_ (7.8125-15.625 Hz), *D*_5_ (3.90625-7.8125 Hz), and *A*_5_ (0-3.90625 Hz) are reserved, as depicted in Fig 4.

### Time-frequency Statistical Feature Extraction

The wavelet components at various levels encompass plentiful time-frequency information, but not all of which are distinctive and pertinent to the task [6]. To filter out non-significant information, a statistical extractor is employed to capture six distinct statistical parameters from each selected wavelet coefficient, as shown in Fig 4. These parameters include mean (*µ*_*ℓq*_), mean absolute deviation (*ρ*_*ℓq*_), standard deviation (*σ*_*ℓq*_), mean absolute value (*m*_*ℓq*_), skewness (*γ*_*ℓq*_), and kurtosis (*κ*_*ℓq*_), similar to the works [16, 20]. The mathematical formulas for these parameters are given as follows:

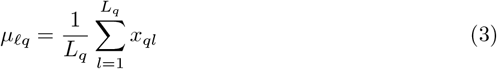

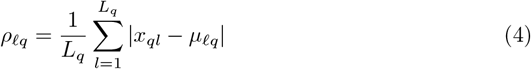

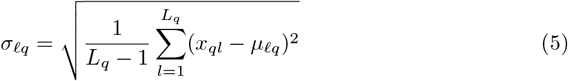

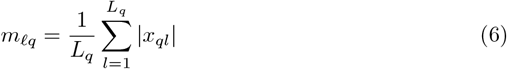

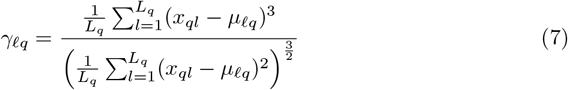

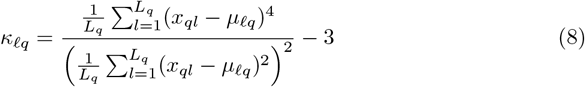

where *ℓ* (*ℓ* = 1,2,…, ℒ) denotes the *ℓ*-th segment of EEG signals, *q* (*q* = 1, 2, 3, 4) represents the *q*-th coefficient belonging to the set {*D*_3_, *D*_4_, *D*_5_, *A*_5_}. *L*_*q*_ is the length of the *q*-th coefficient, and *x*_*ql*_ denotes the *l*-th data point of the *q*-th coefficient. Thus, the mean value serves as a metric for assessing the signal frequency distribution, while the standard deviation and mean absolute deviation quantify variations within the frequency distribution. The mean absolute value gauging the overall amplitude magnitude. Skewness denotes the degree of distortion, and kurtosis characterizes the peakedness of the distribution curve. In this manner, each EEG sample can be transformed into a statistical feature matrix 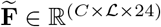:

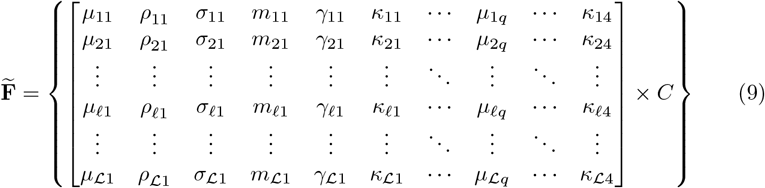

where *C* denotes the count of channels in each EEG sample, ℒ and denotes the count of EEG segments per sample. Notably, for the sake of subsequent analysis, designate the resulting time-frequency feature matrix derived from the *i*-th EEG train trial belonging to class *k*(*k* = 1, 2) as **F**_*ik*_ ∈ ℝ^(*C×ℒ×*24)^. After that, due to the large fluctuations in EEG voltage values, feature-wise Z-score normalization is utilized.

### Spatial Feature Extraction

EEG signals have rich feature expressions in the spatial domain, which has been proven to be an important factor in improving the performance of other EEG classification tasks, such as motor imagery [1, 38, 39], seizure diagnosis [4] and Parkinson’s disease detection [9, 10]. For example, there exist distinctive spatial responses in brain neuro-physiological signals between depression patients and healthy controls [7]. However, most existing EEG pathology studies may not consider spatial information, resulting in limited classification performance. Consequently, this work attempts to mine this important information and integrate it with time-frequency information, aiming to mutually compensate for information deficiencies.

As a renowned supervised spatial feature extraction algorithm, CSP has the ability to capture and augment the spatial distribution information of distinct classes within multi-channel EEG by selecting or weighing the contributions of different spatial regions. Moreover, it has various advantages, such as computational simplicity and powerful dimensionality reduction capability. However, the low SNR nature of EEG signals would compromise the efficacy of CSP, due to its susceptibility to noise [1, 38]. Therefore, in this paper, we implemented CSP on the extracted time-frequency features, excluding noisy information and unrelated temporal and frequency ranges, thereby enhancing the robustness and efficiency of EEG classification. The detailed procedure is as follows.

Firstly, based on the time-frequency features in each EEG test trial for *k*-th category, namely **F**_*ik*_, the averaged normalized covariance matrix 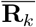 per class is calculated by:

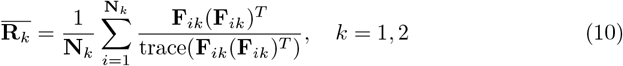

where **N**_*k*_ stands for the sum of trials in *k*-th category, (**F**_*ik*_)^*T*^ represents the transpose of **F**_*ik*_, and the term trace(·) stands for the sum of elements on the diagonal. Thereby, the mixed space covariance matrix **R** can be obtained, and its positive definite nature allows for eigendecomposition through the singular value decomposition theorem :

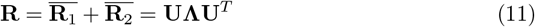

where **U** represents the eigenvector matrix, and **Λ** represents the diagonal matrix with eigenvalues arranged in descending order. Then the whitening transformation matrix **P** can be computed by 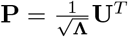, which is applied to concurrently diagonalize the average covariance matrix for two classes:

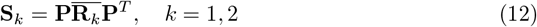

where **S**_*k*_ denotes *k*-th class covariance matrices. **S**_1_ and **S**_2_ share an eigenvector **B**, which can be written as:

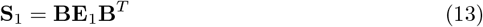

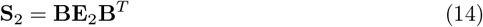

where **E**_1_ and **E**_2_ are eigenvector matrices of **S**_1_ and **S**_2_ respectively. The sum of two eigenvalues from each class of EEG data always results in the identity matrix.

Therefore, **S**_1_ corresponds to the eigenvector with the largest eigenvalue, while **S**_2_ corresponds to the smallest one, and vice versa. Furthermore, the spatial filter **W** can be obtained through a linear transformation, as given by:

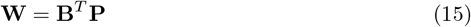

Then input matrices **F**_*i*1_ and **F**_*i*2_ are projected into a low-dimensional space using the spatial filter **W**:

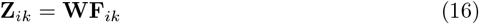

Finally, the features *f*_*ik*_ are normalized by:

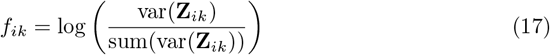

where var(·) and sum(·) are functions that solve for the variance and calculated sum respectively. Through this process, the latent spatial information was converted into apparent low-dimensional eigenvectors with optimal separability, thereby facilitating subsequent classification task.

### Multi-domain Feature Fusion

Both single-domain and dual-domain analyses face the challenge of achieving effective EEG decoding, as features specific to the time, frequency, and spatial domains can only capture EEG signals from their respective perspectives. Multi-domain fusion strategy is a solution to address this challenge as it is proven to be effective in many other EEG analysis tasks [1, 39]. The key motivator behind this strategy is its effectiveness in compensating for the lack of complementarity among heterogeneous characteristics from different domains [1, 30]. Nevertheless, a straightforward feature fusion has the hidden danger of high-dimensional feature space. Specifically, according to the preprocessing applied to the TUH Abnormal EEG dataset, it can be inferred that, for each sample, there is a total of 21 channels and 100 segments respectively. Consequently, the time-frequency feature dimension per sample will then be 21*×* 100*×* 24 = 50400 according to Eq (9). Compared to the spatial feature dimension, which is only 8, it is clear that the former’s dimensionality is extremely high. Such a high dimensionality may give rise to increased model complexity, substantial computational costs, and other issues. As a result, it is necessary to decrease the dimensionality of time-frequency feature space.

Feature aggregation is an excellent countermeasure to condense the extracted features, without compromising feature quality, through the aid of the aggregation function [20]. Therefore, we adopt three Hjorth parameters (i.e., activity, mobility, and complexity) to aggregate each time-frequency statistical feature in every EEG sample from multiple views, as present in Fig 5, owing to their robustness, superior computational efficiency, and the strength of inter-class separation and intra-class aggregation [40]. Take mean for an example:

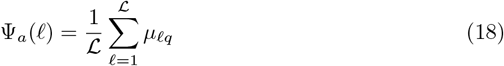

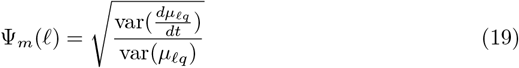

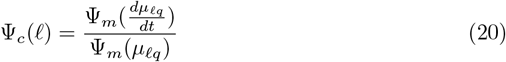

where, Ψ_*a*_(*ℓ*), Ψ_*m*_(*ℓ*), and Ψ_*c*_(*ℓ*) denote the three feature aggregation operations using activity, mobility, and complexity functions, respectively. Thus, time-frequency features per sample are transformed into three *C×* 24 dimensional feature matrices. Then, the aggregated features are fused with spatial features to provide a more comprehensive signal representation in a relatively low-dimensional space. In addition, considering the pronounced correlation between the patient’s age and EEG signals [3, 17], we incorporated it into the feature vector.

**Figure 5.**
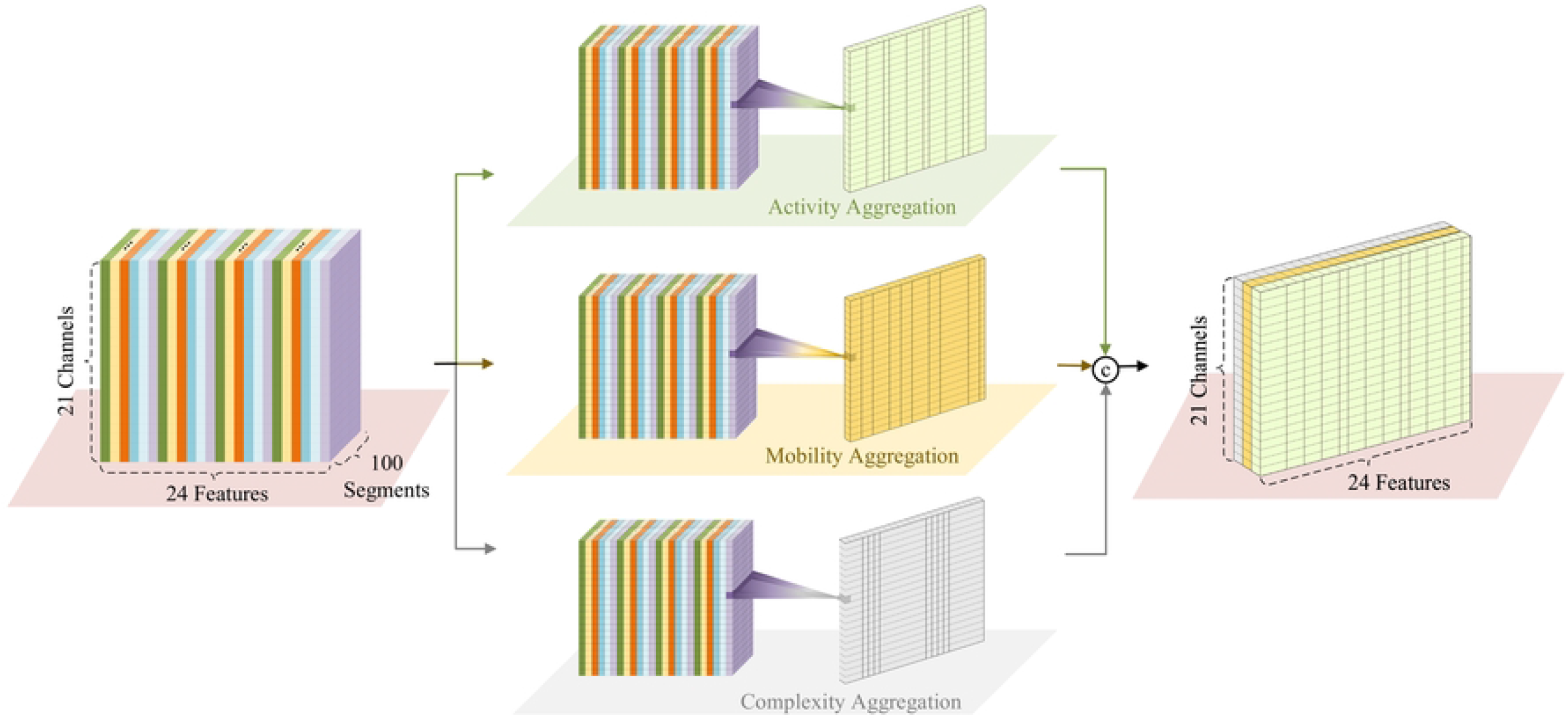
Process of feature aggregation.

### Statistical Significance Analysis

Reducing the fused feature space helps attain a set of features with the best class discrimination ability. However, this process poses a significant challenge for EEG pathology detection. Feature selection, a crucial aspect of this endeavor, eliminates redundant and irrelevant features to reduce the computational burden. While various methods (e.g., neighborhood component analysis [2], ReliefF [1], etc.) have been attempted, the statistical significance analysis-based feature selection has received little attention in EEG pathology detection research. It is noteworthy that as a robust, interpretable, and efficient method, the statistical significance analysis can capture complex relationships between features [41]. Therefore, this study applied this method to select highly distinguishable features.

Given the non-normal distribution of features, the non-parametric Kruskal-Wallis test [42] is utilized to examine features with the most significant statistical impact in classification. The Kruskal-Wallis test formulates the null hypothesis (*H*_0_) that no statistically significant difference exists between independent feature groups, while the alternate hypothesis (*H*_1_) hypothesizes a difference between them. Highly distinguishable features are selected by testing the null hypothesis. In particular, features are ranked according to their discriminative power, and the *p*-value of each feature is computed and compared with the level of significance *α*. Here, *α* is set to be in this work, reflecting a 99% bootstrap Confidence Interval level. If *p*≤ *α, H*_1_ is accepted, indicating that these features are significant for discrimination, and retained. Conversely, when *p > α*, we reject *H*_0_ and discard these insignificant features. Furthermore, a smaller *p*-value implies a more important feature for the given task. Finally, all features with the lower *p*-values (*p <* 0.01) are used to construct a low-dimensional feature matrix and then presented to the classifier.

### Classification

After feature engineering, the last phase is to design a classifier for accurately determining EEG classes. As a multi-classifier ensemble algorithm, GBDT has some advantages, including adaptability to various data distributions and varied feature types, as well as the capability to handle complex nonlinear relationships, contributing to its high predictive accuracy across a wide range of applications [4, 20, 24]. In light of the nonlinear and non-Gaussian nature of EEG data, we employed GBDT’s recent and prominent implementations, namely CatBoost, XGBoost, and LightGBM, for discriminating between normal and abnormal EEG signals and evaluating the proposed feature engineering, as detailed in Section. The results indicate that CatBoost achieves the best performance among these three classifiers (refer to Fig 6). Consequently, we integrated the proposed feature engineering with CatBoost for comparison against several existing approaches.

**Figure 6.**
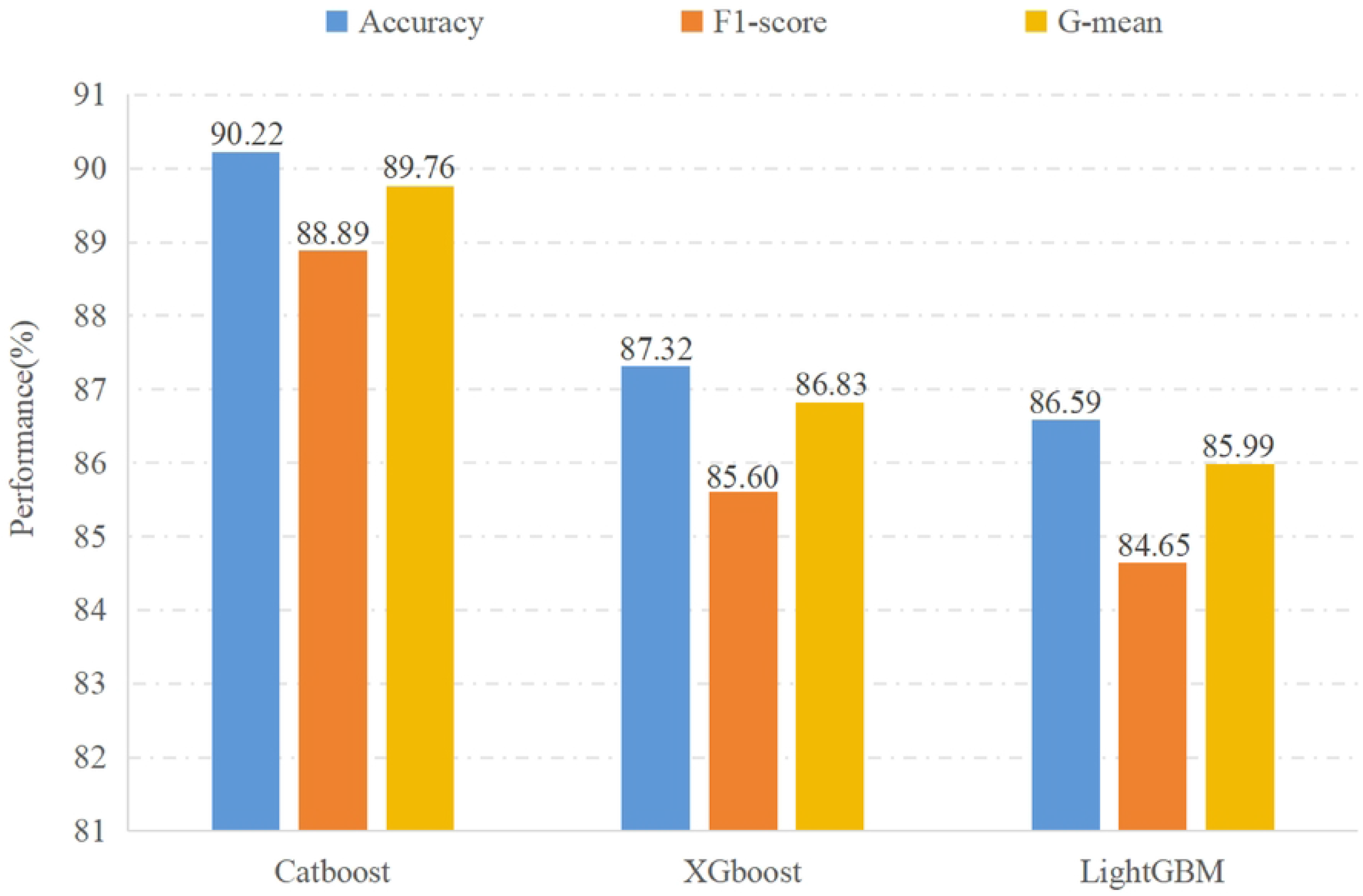
Performance of three classifiers.

## Results and Discussion

In this section, we begin by presenting the experimental setup and evaluation metrics. Subsequently, we undertake a comprehensive evaluation, contrasting the proposed methods with several state-of-the-art deep learning and feature-based methods. Finally, two ablation studies are performed to validate the impact of key components in the proposal. This method was constructed using the scikit-learn library [43], scripted in Python3, on an Ubuntu 18.04.6 LTS. All experiments were executed on a workstation equipped with an AMD Ryzen Threadripper 3970 32-Core Processor and 128 GB RAM.

### Experimental Setup

In order to evaluate the efficacy of the proposed framework, extensive experiments were conducted on the TUH Abnormal EEG database. Raw EEG data was preprocessed through standard 21-channel selection, downsampling to 250Hz, and 5 s non-overlapping segmentation. In the feature extraction stage, six wavelet-based time-frequency features, i.e., mean, mean deviation, standard deviation, mean absolute value, skewness, and kurtosis were extracted, as elaborated in Section. Spatial features were derived from the constructed time-frequency feature space via CSP with eight decomposition components. Simultaneously, activity, mobility, and complexity mapped the time-frequency features into a lower-dimensional feature space, which is subsequently integrated with spatial and age features. To mitigate redundant and irrelevant features, we chose highly discriminative features with *p*-values below 0.01 and fed them into CatBoost, XGBoost, and LightGBM. The hyper-parameters of these classifiers were fine-tuned as follows: 1080 estimator counts, maximum depths of 4, 6, and 4, and learning rates of 0.03, 0.02, and 0.04, respectively; other hyper-parameters were kept at default values. Consistent with established practices [20, 32], the system was separately trained and tested on mutually independent training and testing sets, both sourced from TUH Abnormal EEG.

To evaluate the effectiveness of pathological EEG detection, three common evaluation metrics, namely accuracy, F1-score, and G-mean, are utilized. Accuracy is an intuitive and standard criterion that represents the proportion of correctly classified instances relative to all instances. Besides, due to the unbalanced class distributions in the training set (refer to Table 1), accuracy is insufficient for comprehensive assessment [16]. Hence, we incorporated both the F1-score and G-mean, each offering a balanced view of the model’s performance, to optimize our evaluation. F1-score serves as a metric that harmoniously combines precision and recall through the computation of their harmonic mean, while the G-mean represents the exact geometric mean of the recalls for both positive and negative classes.

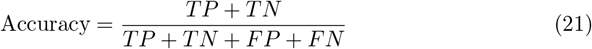

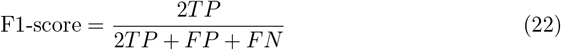

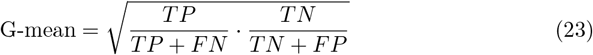

where, TP (True Positive) and TN (True Negative) denote the accurate identification counts of positive and negative instances, respectively. FP (False Positive) represents the incorrect assignment of normal instances to the abnormal class, while FN (False Negative) indicates the misclassification of abnormal instances. The confusion matrix, as present in Table 2, encompasses these four metrics.

**Table 2.**
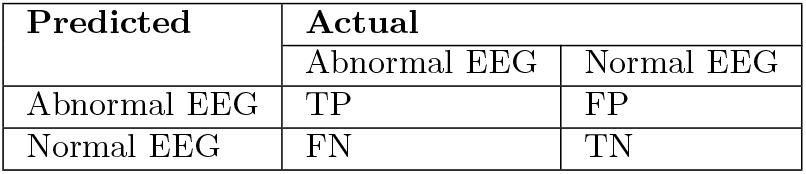
Confusion matrix of EEG detection.

| Predicted | Actual |  |
| --- | --- | --- |
|  | Abnormal EEG | Normal EEG |
| Abnormal EEG | TP | FP |
| Normal EEG | FN | TN |

### Performance Testing on the Extracted Features with Different Classifiers

To evaluate the effectiveness and broad applicability of the proposed feature engineering, this study employed three advanced ensemble learning classifiers: CatBoost, XGBoost, and LightGBM, for categorizing the extracted EEG pathological features. The results obtained using each classifier are presented in Fig 6.

Fig 6 indicates that among these ensemble learning classifiers, the framework coupled with CatBoost exhibited superior performance in distinguishing between non-pathological and pathological EEGs with an accuracy rate, F1-score, and G-mean of 90.22%, 88.89%, and 89.76% respectively. It is noteworthy that the accuracy of this method exceeds the clinical diagnostic standard, which typically mandates a reliable system to attain an accuracy of 90% for the diagnosis of abnormal EEG [11], indicating its medical reliability and practical utility. When applying XGBoost and LightGBM, both frameworks demonstrated commendable classification accuracy, achieving 87.32% and 86.59% respectively. These favorable performances are also evident in the F1-score and G-mean, confirming the extensive applicability and stability of the proposed feature engineering across different machine-learning frameworks. In a word, our feature engineering attains satisfactory classification performance, with accuracy consistently exceeding 86.5% in all cases, robustly supporting the feasibility of multi-domain feature fusion and two-step dimension reduction. Additionally, the outstanding performance of the CatBoost-based approach prompts us to utilize it in subsequent experiments.

The confusion matrices for EEG pathology detection using the above three frameworks on the TUH Abnormal EEG Database are presented in Fig 7. We can observe that, based on the input features, CatBoost misclassified 18 out of 126 abnormal instances and 9 out of 150 normal instances, resulting in a higher false negative rate of 14.29% than the false positive rate of 6%. Similar phenomena, where the model excels in accurately identifying normal EEGs, are evident in the other two classifiers. These findings substantiate that all three classifiers exhibit high sensitivity to normal class, which is especially pivotal in automated diagnosis [20]. Interestingly, this result is in alignment with previous research studies [11, 32]. The reason could possibly be the unequal distribution of different classes in the training set (refer to Table 1), specifically the larger quantity of normal EEG, which could induce a bias towards this class in the method.

**Figure 7.**
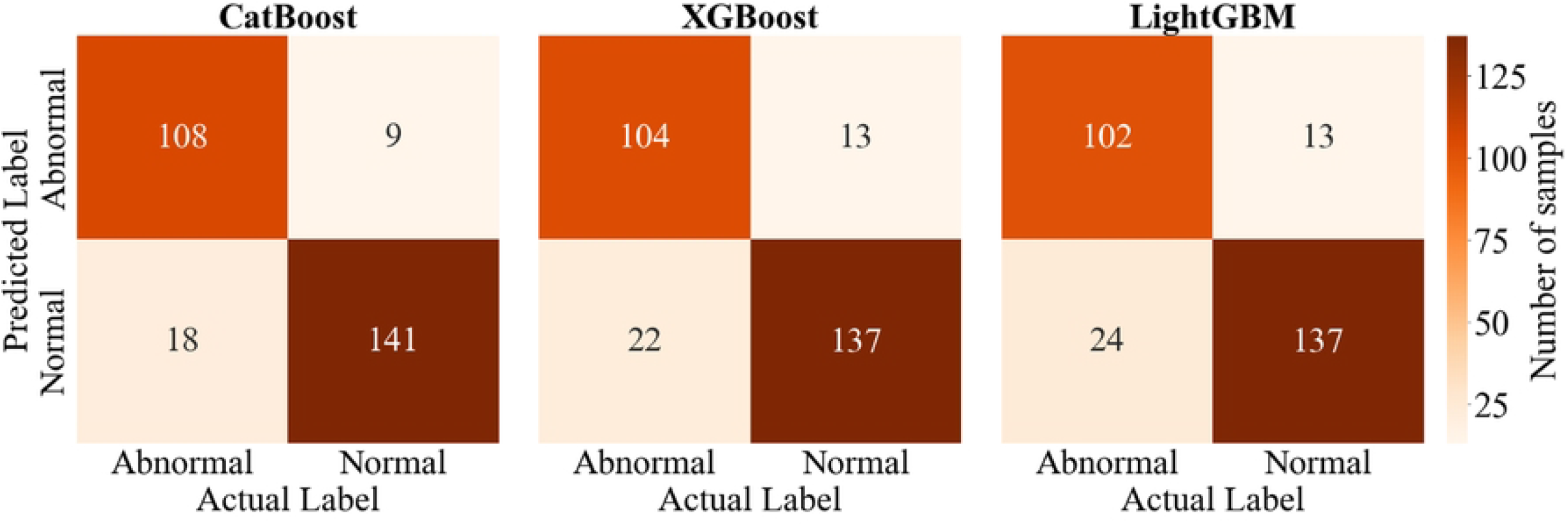
Confusion matrices of three classifiers.

### Comparative Analysis with Existing State-of-the-art Methods

To demonstrate the superiority of our methodology, a comparison against several representative deep learning and feature-based approaches was carried out on the TUH Abnormal EEG Corpus, as presented in Table 3.

**Table 3.** Comparison of the classification results obtained by different EEG pathology diagnosis approaches on the real-world EEG abnormal dataset.

| Category | Reference | Features | Classifier | Accuracy(%) | F1-score (%) | G-mean (%) |
| --- | --- | --- | --- | --- | --- | --- |
| Deep Learning Approach | [31] | - | BD-Deep4 | 85.40 | 82.52 | 84.08 |
|  | [22]<sup>*</sup> | - | AlexNet + MLP | 89.13 | 87.06 | 88.02 |
|  | [32]<sup>*</sup> | - | AlexNet + SVM | 87.32 | 84.97 | 86.24 |
|  | [33] | - | WaveNet-LSTM | 88.76 | 88.32 | 88.39 |
| Feature-based Approach | [11] | DWT, CWT, DFT | HT+RG | 85.86 | 83.4 | 85.19 |
|  | [13] | Statistical features + spectral power | LSTM + Attention | 79.05 | 79.00 | 79.00 |
|  | [20] | WPD | CatBoost | 87.68 | 86.06 | 87.24 |
|  | [2] | Multilevel DWT | KNN | 87.68 | 86.07 | 87.24 |
|  | [24] | WPD | CatBoost | 89.13 | 87.60 | 88.60 |
|  | Ours | DWT + CSP | CatBoost | **90.22** | **88.89** | **89.76** |

\*Additional closed-source data is employed for training. The best results are highlighted in bold.

Table 3 illustrates that our method exhibits superior performance than other state-of-the-art frameworks. Through comprehensive analysis, we can obtain the following conclusions: (i) In contrast to four deep learning methodologies, the proposed framework consistently achieves the highest accuracy, F1-score, and G-mean. Specifically, compared against two transfer learning methodologies [22, 32], both of which were additionally trained on undisclosed EEG data, our approach exhibits notable improvements of 1.09% and 2.90% in accuracy, 1.83% and 3.92% in F1-score, as well as 1.74% and 3.52% in G-mean. This phenomenon can be ascribed to the dependency of deep learning methods on complex end-to-end structures with numerous parameters and a substantial amount of training samples, causing a potential risk of overfitting and increased computational resources. Moreover, the black-box nature of deep learning approaches is another factor impacting their performance. The opaque internal decision logic of these models results in poor interpretability which is crucial in medical applications to ensure diagnostic reliability and safety. Besides, the oversight of subtle spatial information restricts the upper limitation of these methodologies’ performance. In contrast, our study turns to a simple and reliable multi-domain feature-based approach, which achieves higher EEG detection accuracy with lower computational resource consumption.

(ii) Also, our framework outperforms five advanced feature-based methods. Specifically, it surpasses two methodologies [2, 11] that employ the same time-frequency feature extraction technique as ours, namely DWT. This advancement can be attributed to three key advantages. Firstly, the proposed spatial feature representation compensates for the lack of time-frequency features. The second factor is the complementary information from multi-domain fusion features through comprehensive multi-feature fusion. Thirdly, a two-step dimension reduction strategy efficiently diminishes data dimensions and selects more representative features for EEG pathological detection. On the other hand, our method outperforms the suboptimal feature-based approach [24] in accuracy, F1-score, and G-mean by 1.09%, 1.29%, and 1.16%, respectively. It is noteworthy that this work employs the same classifier as our framework, further signifying the effectiveness of the proposed feature engineering.

In summary, these results validate the benefits of combining spatial information with time-frequency domain feature extraction and a two-step dimension reduction strategy in EEG modeling. Therefore, this work opted to construct a novel feature-based traditional machine learning methodology coupled with these key components to tackle the challenges in abnormal EEG detection.

### Ablation Study

The multi-view feature aggregation and spatial information learning are two pivotal elements that affect the performance and efficiency of the EEG binary classification. Hence, in this section, two ablation experiments were conducted on the TUH Abnormal EEG dataset to further investigate and evaluate the individual impact of these two components.

### Effect of Multi-view Feature Aggregation

The multi-view feature aggregation mechanism is a practical implement to reduce the dimension of time-frequency feature space and simplify the model complexity. To fully investigate the effect of feature aggregation, we performed an ablation experiment by gradually increasing the aggregate function. Here, four different cases were compared and analyzed, including:

- Case-1: Without feature aggregation, time-frequency, spatial, and age features are directly concatenated into a feature vector and then input into CatBoost.
- Case-2: Time-frequency features are aggregated using activity and then concatenated with spatial and age features.
- Case-3: Time-frequency features are aggregated using activity and mobility simultaneously and then concatenated with spatial and age features.
- Case-4: Time-frequency features are aggregated using activity and complexity simultaneously and then concatenated with spatial and age features.
- Proposed method: Time-frequency features are aggregated using activity, mobility, and complexity simultaneously and then concatenated with spatial and age features.

To ensure experimental fairness, this study utilized a uniform feature set and a unified CatBoost classifier across all cases. Additionally, the execution time covering feature learning and classification was taken into account for assessing the effect of the mechanism in classification efficiency. Table 4 presents the outcomes of five cases. A comparative analysis reveals several key insights:

**Table 4.** Ablation study of multi-view feature aggregation.

| Case | Feature aggregation |  |  | Evaluation metrics |  |  |  |
| --- | --- | --- | --- | --- | --- | --- | --- |
|  | activity | mobility | complexity | Accuracy (%) | F1-score (%) | G-mean (%) | Execution time (s) |
| Case-1 |  |  |  | 81.52 | 78.66 | 80.71 | 194.43 |
| Case-2 | ✓ |  |  | 86.23 | 84.55 | 85.87 | 53.65 |
| Case-3 | ✓ | ✓ |  | 88.04 | 86.53 | 87.66 | 55.31 |
| Case-4 | ✓ |  | ✓ | 86.96 | 85.60 | 86.77 | 55.38 |
| Proposed method | ✓ | ✓ | ✓ | **90.22** | **88.89** | **89.76** | 60.80 |

The best results are highlighted using bold italics.

1. Even without utilizing feature aggregation, Case-1 exhibits a classification accuracy of 81.52%, surpassing the comparative method [13]. This improvement is consistent across the other two indicators, revealing that the proposed spatial information learning and the multi-domain feature fusion can improve the accuracy of EEG decoding.
2. It is evident that the performance of feature aggregation methods demonstrates a monotonic increase as the aggregation function increments. Specifically, in contrast to Case-1, Case-2 adopting one Hjorth parameter (i.e., activity) yields an accuracy improvement of 4.71% and a substantial time consumption reduction of 140.78 seconds. This phenomenon can be attributed to that feature aggregation effectively compresses high-dimensional features, thereby improving feature recognizability, reducing computational overhead, and enhancing classification performance. Compared to

Case-2, Case-3 and Case-4 incorporating a second Hjorth parameter, namely mobility and complexity respectively, lead to improvements in accuracy by 1.81% and 0.73%, F1-scores of 1.98% and 1.05%, along with G-mean of 1.79% and 0.9%. Further analysis revealed that when all Hjorth parameters were employed, the proposed method achieved the peak values across accuracy, F1-score, and G-mean. These results suggest that each Hjorth parameter provides unique insights regarding amplitude, frequency, and other waveform characteristics so that multi-view information integration can comprehensively characterize the time-frequency features, thereby enhancing the power of aggregated features for discriminating EEG signals. Although our method costs a slightly higher time cost compared to Cases -2, -3, and -4, its optimal classification performance justifies the time investment. Meanwhile, in comparison to Case-1, our method reduced execution time by 133.63 seconds and significantly increased the accuracy rate of EEG signal detection by 8.70%.

In summary, these experimental results substantiate that this multi-view feature aggregation mechanism effectively condenses information by diverse viewpoints to a lower-dimensional and more discriminative information representation. Therefore, this mechanism not only enables the method to efficiently deal with high-dimensional data but also accelerates the execution speed of EEG classification.

### Effect of Spatial Information Learning

In recent years, the majority of feature engineering focuses on exploring time, frequency, and time-frequency domains, with scant attention given to the spatial domain, which is pivotal for improving the precision of EEG pathology detection. In this subsection, to validate the vital effect of spatial information, we compare and analyze two distinct cases: (1) the resultant feature set contains the spatial domain features and (2) the spatial features are left out before classification. To mitigate the inherent impact of the classifier on comparisons, three ensemble learning classifiers, consistent with those in Section, were used to handle the feature sets. The classification performances of these two scenarios are illustrated respectively in Figs 6 and 8.

**Figure 8.**
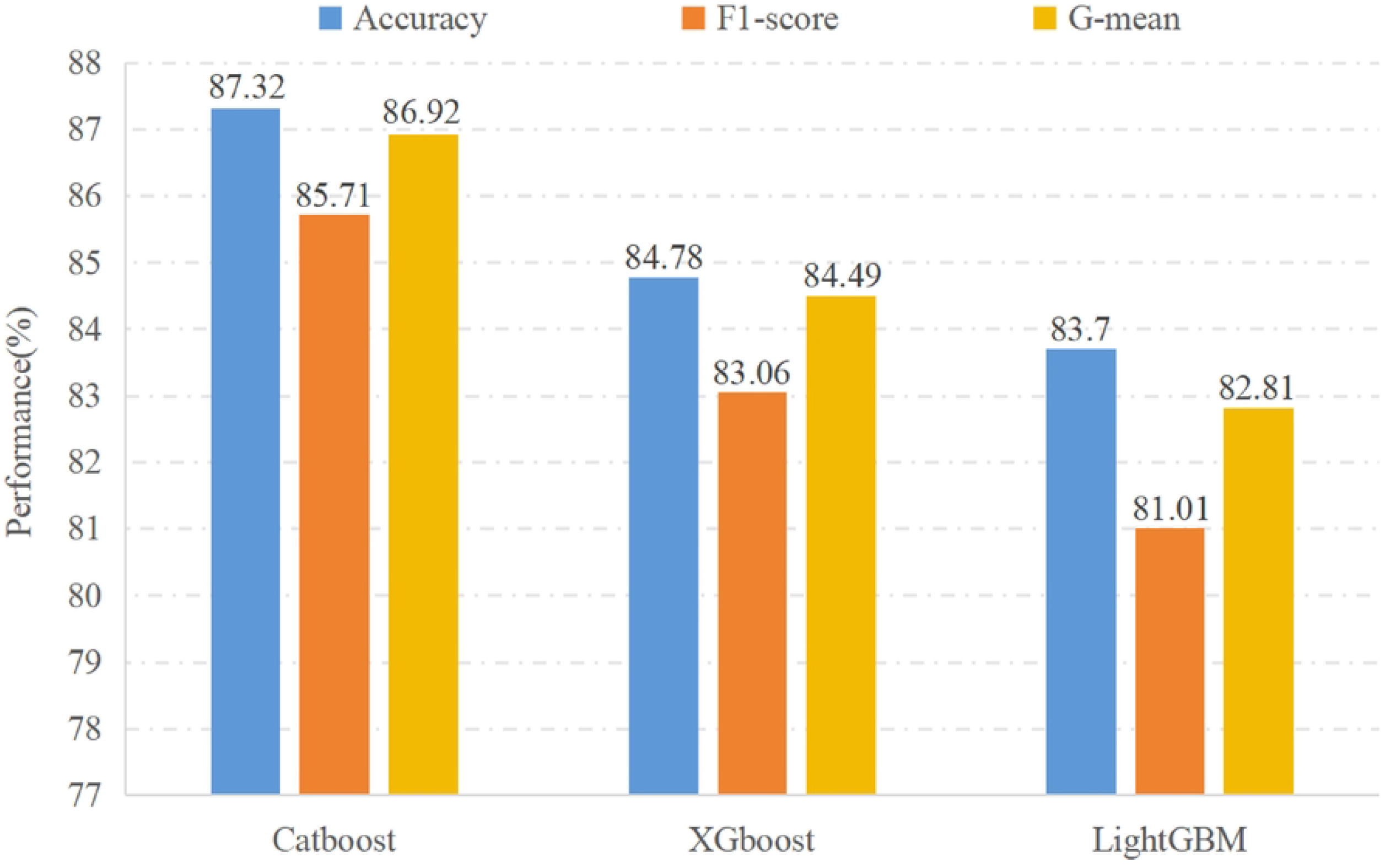
Performance of three classifiers (without spatial feature

Fig 8 depicts the results of Case-1 exclusively learning the time-frequency features. It is evident that CatBoost-based approach achieves accuracy, F1-score, and G-mean of 87.32%, 85.71%, and 86.92% respectively, surpassing most contrast algorithms [11, 13, 31, 32]. This remarkable performance provides supporting evidence for the excellence of the proposed time-frequency feature representation. If these results are further compared against the results in Fig 6, a substantial improvement in classification performance can be observed across all three approaches, attributable to the inclusion of time-frequency spatial information. Taking Catboost as an example, there is a prominent increase in accuracy, F1-score, and G-mean by 2.90%, 3.18%, and 2.84%, respectively. It indicates that the latent spatial information, which carries the important spatial distribution of different classes within the multi-channel EEG signals, effectively compensates for the information gap in the time-frequency feature.

Interestingly, this result aligns with [5] affirming that the combination of time, frequency, and spatial features effectively enhances seizure detection.

## Conclusion

In this work, we designed an automatic system for detecting EEG abnormalities, with the objective of aiding in the treatment of EEG-related neurological conditions. This automatic system adopts an innovative feature-based architecture to comprehensively mine EEG signals and enrich the signal representation, thereby improving the accuracy of EEG decoding. On one hand, a multi-domain feature fusion model is designed to fully account for complementary information among multi-domain features. On the other hand, an innovative two-step dimensionality reduction strategy is implemented to improve the capability of feature representation, laying a foundation for the classification. The comparative experiments were conducted on the TUH Abnormal EEG dataset, where the proposed methodology significantly outperforms the cutting-edge representative deep learning and feature-based baselines, substantiating the efficacy and feasibility of our method. Besides, a series of ablation assessments signified the important effect of spatial features and dimension reduction in the EEG diagnosis.

Despite the noticeable performance, there are still several avenues for future exploration. One interesting direction would be the development of an adaptive feature selection method capable of dynamically identifying discriminative features without a predefined threshold for examining features, providing an enhanced alternative to statistical testing-based selection. Another is to augment model effectiveness through the optimization of brain electrical channels.

## Author Contributions

**Conceptualization:** Shimiao Chen, Xiangzeng Kong.

**Data curation:** Shimiao Chen, Dong Huang, Xinyue Liu.

**Formal analysis:** Shimiao Chen, Xinyue Liu.

**Funding acquisition:** Xiangzeng Kong, Tingting Zhang.

**Investigation:** Shimiao Chen, Xinyue Liu, Jianjun Chen.

**Methodology:** Shimiao Chen, Xiangzeng Kong.

**Project administration:** Xiangzeng Kong, Tingting Zhang.

**Resources:** Xiangzeng Kong, Tingting Zhang.

**Software:** Shimiao Chen, Dong Huang, Xinyue Liu.

**Supervision:** Xiangzeng Kong, Tingting Zhang.

**Validation:** Shimiao Chen, Xinyue Liu, Jianjun Chen, Xiangzeng Kong.

**Visualization:** Xinyue Liu, Jianjun Chen.

**Writing - original draft:** Shimiao Chen, Jianjun Chen.

**Writing - review & editing:** Shimiao Chen, Jianjun Chen.

## References

1. Zheng L, Feng W, Ma Y, Lian P, Xiao Y, Yi Z, et al. Ensemble learning method based on temporal, spatial features with multi-scale filter banks for motor imagery EEG classification. Biomed Signal Process Control. 2022;76:103634. doi:10.1016/j.bspc.2022.103634.

2. Tasci I, Tasci B, Barua PD, Dogan S, Tuncer T, Palmer EE, et al. Epilepsy detection in 121 patient populations using hypercube pattern from EEG signals. Inf Fusion. 2023;96:252–268. doi:10.1016/j.inffus.2023.03.022.

3. Khan HA, Ul Ain R, Kamboh AM, Butt HT, Shafait S, Alamgir W, et al. The NMT Scalp EEG Dataset: An Open-Source Annotated Dataset of Healthy and Pathological EEG Recordings for Predictive Modeling. Front Neurosci. 2022;15. doi:10.3389/fnins.2021.755817.

4. Wu D, Li J, Dong F, Liu J, Jiang L, Cao J, et al. Classification of seizure types based on multi-class specific bands common spatial pattern and penalized ensemble model. Biomed Signal Process Control. 2023;79:104118. doi:10.1016/j.bspc.2022.104118.

5. Huang H, Chen P, Wen J, Lu X, Zhang N. Multiband seizure type classification based on 3D convolution with attention mechanisms. Comput Biol Med. 2023;166:107517. doi:10.1016/j.compbiomed.2023.107517.

6. Shen M, Wen P, Song B, Li Y. An EEG based real-time epilepsy seizure detection approach using discrete wavelet transform and machine learning methods. Biomed Signal Process Control. 2022;77:103820. doi:10.1016/j.bspc.2022.103820.

7. Jiang C, Li Y, Tang Y, Guan C. Enhancing EEG-Based Classification of Depression Patients Using Spatial Information. IEEE Trans Neural Syst Rehabil Eng. 2021;29:566–575. doi:10.1109/TNSRE.2021.3059429.

8. Minkowski L, Mai KV, Gurve D. Feature Extraction to Identify Depression and Anxiety Based on EEG. In: 2021 43rd Annual International Conference of the IEEE Engineering in Medicine and Biology Society (EMBC); 2021. p. 6322–6325.

9. Aljalal M, Aldosari SA, AlSharabi K, Abdurraqeeb AM, Alturki FA. Parkinson’s Disease Detection from Resting-State EEG Signals Using Common Spatial Pattern, Entropy, and Machine Learning Techniques. Diagn. 2022;12(5). doi:10.3390/diagnostics12051033.

10. Chang H, Liu B, Zong Y, Lu C, Wang X. EEG-Based Parkinson’s Disease Recognition via Attention-Based Sparse Graph Convolutional Neural Network. IEEE J Biomed Health Inform. 2023;27(11):5216–5224. doi:10.1109/JBHI.2023.3292452.

11. Gemein LAW, Schirrmeister RT, Chrabaszcz P, Wilson D, Boedecker J, Schulze-Bonhage A, et al. Machine-learning-based diagnostics of EEG pathology. NeuroImage. 2020;220. doi:10.1016/j.neuroimage.2020.117021.

12. Sharma M, Patel S, Acharya UR. Automated detection of abnormal EEG signals using localized wavelet filter banks. Pattern Recognit Lett. 2020;133:188–194. doi:10.1016/j.patrec.2020.03.009.

13. Cisotto G, Zanga A, Chlebus J, Zoppis I, Manzoni S, Markowska-Kaczmar U. Comparison of attention-based deep learning models for eeg classification. arXiv preprint 201201074. 2020;.

14. Yildirim Ö, Baloglu UB, Acharya UR. A deep convolutional neural network model for automated identification of abnormal EEG signals. Neur Comput Appl. 2020;32:15857–15868. doi:10.1007/s00521-018-3889-z.

15. Peh WY, Thomas J, Bagheri E, Chaudhari R, Karia S, Rathakrishnan R, et al. Multi-Center Validation Study of Automated Classification of Pathological Slowing in Adult Scalp Electroencephalograms Via Frequency Features. Int J Neural Syst. 2021;31(06):2150016. doi:10.1142/S0129065721500167.

16. Bajpai R, Yuvaraj R, Prince AA. Automated EEG pathology detection based on different convolutional neural network models: Deep learning approach. Comput Biol Med. 2021;133:104434. doi:10.1016/j.compbiomed.2021.104434.

17. Kiessner AK, Schirrmeister RT, Gemein LAW, Boedecker J, Ball T. An extended clinical EEG dataset with 15,300 automatically labelled recordings for pathology decoding. NeuroImage Clin. 2023;39:103482. doi:10.1016/j.nicl.2023.103482.

18. López S, Suarez G, Jungreis D, Obeid I, Picone J. Automated identification of abnormal adult EEGs. In: 2015 IEEE Signal Processing in Medicine and Biology Symposium (SPMB); 2015. p. 1–5.

19. Roy S, Kiral-Kornek I, Harrer S. ChronoNet: A deep recurrent neural network for abnormal EEG identification. In: Artificial Intelligence in Medicine: 17th Conference on Artificial Intelligence in Medicine, AIME 2019, Poznan, Poland, June 26–29, 2019, Proceedings 17. Springer; 2019. p. 47–56.

20. Albaqami H, Hassan GM, Subasi A, Datta A. Automatic detection of abnormal EEG signals using wavelet feature extraction and gradient boosting decision tree. Biomed Signal Process Control. 2021;70:102957. doi:10.1016/j.bspc.2021.102957.

21. Ding Y, Meng Y, Wang L. Study on real-time prediction method of seizures based on yolov3 for EEG spike wave detection. In: Proceedings of the 3rd International Conference on Computer Science and Application Engineering; 2019. p. 1–7.

22. Alhussein M, Muhammad G, Hossain MS. EEG Pathology Detection Based on Deep Learning. IEEE Access. 2019;7:27781–27788. doi:10.1109/ACCESS.2019.2901672.

23. Blanco S, Garcia H, Quiroga RQ, Romanelli L, Rosso OA. Stationarity of the EEG series. IEEE Eng Med Biol Mag. 1995;14(4):395–399. doi:10.1109/51.395321.

24. Zhong Y, Wei H, Chen L, Wu T. Automated EEG Pathology Detection Based on Significant Feature Extraction and Selection. Math. 2023;11(7). doi:10.3390/math11071619.

25. Singh R, Ahmed T, Kumar Singh A, Chanak P, Singh SK. SeizSClas: An Efficient and Secure Internet-of-Things-Based EEG Classifier. IEEE IoT J. 2021;8(8):6214–6221. doi:10.1109/JIOT.2020.3030821.

26. Pradhan BK, Jarzebski M, Gramza-Michalowska A, Pal K. Automated Detection of Caffeinated Coffee-Induced Short-Term Effects on ECG Signals Using EMD, DWT, and WPD. Nutrients. 2022;14(4). doi:10.3390/nu14040885.

27. Kohad N, Ramesh R, Roy R, Irrinki S, S N. Segment Based Abnormality Detection in EEG Recordings. In: 2022 2nd International Conference on Intelligent Technologies (CONIT); 2022. p. 1–8.

28. Tsipouras MG. Spectral information of EEG signals with respect to epilepsy classification. EURASIP J Adv Signal Process. 2019;2019. doi:10.1186/s13634-019-0606-8.

29. Wei S, Gao J, Yang Y, Xiong N, Zhang J, Song J, et al. Analysis of Weight-Directed Functional Brain Networks in the Deception State Based on EEG Signal. IEEE J Biomed Health Inform. 2023;27(10):4736–4747. doi:10.1109/JBHI.2023.3295892.

30. Lin R, Dong C, Ma P, Ma S, Chen X, Liu H. A Fused Multidimensional EEG Classification Method Based on an Extreme Tree Feature Selection. Comput Intell Neurosci. 2022;2022. doi:10.1155/2022/7609196.

31. Schirrmeister RT, Springenberg JT, Fiederer LDJ, Glasstetter M, Eggensperger K, Tangermann M, et al. Deep learning with convolutional neural networks for EEG decoding and visualization. Hum Brain Mapp. 2017;38(11):5391–5420. doi:10.1002/hbm.23730.

32. Amin SU, Hossain MS, Muhammad G, Alhussein M, Rahman MA. Cognitive smart healthcare for pathology detection and monitoring. IEEE Access. 2019;7:10745–10753. doi:10.1109/ACCESS.2019.2891390.

33. Albaqami H, Hassan GM, Datta A. Automatic Detection of Abnormal EEG Signals Using WaveNet and LSTM. Sens. 2023;23(13). doi:10.3390/s23135960.

34. Mohsenvand MN, Izadi MR, Maes P. Contrastive Representation Learning for Electroencephalogram Classification. In: Proceedings of the Machine Learning for Health NeurIPS Workshop. vol. 136 of Proceedings of Machine Learning Research. PMLR; 2020. p. 238–253.

35. Obeid I, Picone J. The Temple University Hospital EEG Data Corpus. Front Neurosci. 2016;10. doi:10.3389/fnins.2016.00196.

36. Jiwani N, Gupta K, Afreen N. Automated Seizure Detection using Theta Band. In: 2022 International Conference on Emerging Smart Computing and Informatics (ESCI); 2022. p. 1–4.

37. Frikha T, Abdennour N, Chaabane F, Ghorbel O, Ayedi R, Shahin OR, et al. Source Localization of EEG Brainwaves Activities via Mother Wavelets Families for SWT Decomposition. J Healthc Eng. 2021;2021. doi:10.1155/2021/9938646.

38. Rithwik P, Benzy VK, Vinod AP. High accuracy decoding of motor imagery directions from EEG-based brain computer interface using filter bank spatially regularised common spatial pattern method. Biomed Signal Process Control. 2022;72:103241. doi:10.1016/j.bspc.2021.103241.

39. Zhang Y, Chen W, Lin CL, Pei Z, Chen J, Chen Z. Boosting-LDA algriothm with multi-domain feature fusion for motor imagery EEG decoding. Biomed Signal Process Control. 2021;70:102983. doi:10.1016/j.bspc.2021.102983.

40. Liang T, Lu H. A Novel Method Based on Multi-Island Genetic Algorithm Improved Variational Mode Decomposition and Multi-Features for Fault Diagnosis of Rolling Bearing. Entropy. 2020;22(9). doi:10.3390/e22090995.

41. Siuly S, Khare SK, Bajaj V, Wang H, Zhang Y. A Computerized Method for Automatic Detection of Schizophrenia Using EEG Signals. IEEE Trans Neural Syst Rehabil Eng. 2020;28(11):2390–2400. doi:10.1109/TNSRE.2020.3022715.

42. Kruskal WH, Wallis WA. Use of Ranks in One-Criterion Variance Analysis. J Am Stat Assoc. 1952;47(260):583–621. doi:10.1080/01621459.1952.10483441.

43. Pedregosa F, Varoquaux G, Gramfort A, Michel V, Thirion B, Grisel O, et al. Scikit-learn: Machine learning in Python. J Mach Learn Res. 2011;12:2825–2830.

